# Functional ultrasound imaging of deep visual cortex in awake non-human primates

**DOI:** 10.1101/551663

**Authors:** Blaize Kévin, Gesnik Marc, Arcizet Fabrice, Ahnine Harry, Ferrari Ulisse, Deffieux Thomas, Pouget Pierre, Chavane Frédéric, Fink Mathias, Sahel José-Alain, Tanter Mickael, Picaud Serge

**Author notes:** BK and GM are Co-first authors. TM and PS are Co-last authors. Contacts: BK, PS.

## Abstract

Deep regions of the brain are not easily accessible to investigation at the mesoscale level in awake animals or humans. We have recently developed functional Ultrasound (fUS) imaging fUS imaging technique to uncover deep hemodynamic functional responses. Applying fUS imaging on two awake non-human primates performing a passive fixation task, we reconstructed their retinotopic maps down to the deep calcarine and lunate sulci on visual areas (V1, V2 and V3). These maps were acquired in a single hour session with very few stimuli presentation. The spatial resolution of the technology is illustrated by mapping of Ocular Dominance (OD) columns within superficial and deep layers of the primary visual cortex. These acquisitions showed that OD selectivity is mostly present in layer IV but with evidence also in layers II/III and V. The fUS imaging technology therefore provides a new mesoscale approach to map brain activities at high spatiotemporal resolution in awake subjects within the whole depth of the cortex.

## Introduction

To understand brain function in normal and pathological conditions, novel technologies are needed to measure and map brain activities in large animal models and patients. For evaluating these technologies, the mammalian visual system offers an ideal system because it contains many visual areas that collectively embrace a large fraction of the cerebral cortex (Felleman and Van Essen, 1991). Furthermore, this system was highly characterized thanks to the possibility for very precise visual stimulations in the visual field. For neuroscientists interested in vision, a fundamental set of objectives is to identify at the mesoscale the overall extent of visual cortex while at the microscale to reveal concepts into the neuronal computations that underlie behavior. Recent advances in neurotechnology have shown insights into the neuronal computations that underlie behavior at the microcircuit level in rodents (Callaway and Luo, 2015; Kerr and Denk, 2008; Macé et al., 2018) or at the macro-circuit level in non-human primates (NHP) using functional Magnetic Resonance Imaging (fMRI) as imaging approach (Tsao et al., 2008; Vanduffel et al., 2001; Yue et al., 2013). However, a combined macro and micro approach to uncover activity maps during behaving tasks is necessary to achieve a system-level understanding of visual information processing.

fMRI in visual cortex studies has given access to low resolution retinotopic maps but ocular dominance bands were too small to be resolved in NHP (Benson et al., 2012; Engel et al., 1994; Henriksson et al., 2012). Optical imaging provides a microscopic resolution in non-human primates using either Intrinsic Optical imaging (Blasdel and Campbell, 2001; Heider et al., 2005; Ramsden et al., 2001; Ts’o et al., 1990; Vanzetta et al., 2004), voltage sensitive dyes imaging (VSDI) (Chemla et al., 2017; Chen et al., 2006; Grinvald et al., 1994; Meirovithz et al., 2010; Muller et al., 2014; Reynaud et al., 2012) or two-photon microscopy using calcium sensors (Murakami et al., 2015; Nauhaus et al., 2016; Zhuang et al., 2017). The main disadvantages of these approaches lie either of being restricted to the cortical surface (optical imaging, VSDI) or otherwise limited in the imaging depth and the restricted field of view (FOV) (two-photon microscopy, up to ~600μm) (Helmchen and Denk, 2005; Oheim et al., 2001). Consequently, these observations were not accessible to the deep cortical folds of the visual cortex within the Calcarine sulcus (Hinds et al., 2008; Stensaas et al., 1974). Recently, the new functional ultrasound (fUS) imaging technique was found to provide a high spatiotemporal resolution (100μm, 1Hz) even in deep structures (up to 1.5cm) (Gesnik et al., 2017; Macé et al., 2011) and applied to map in 3D the visual system of rodents (Gesnik et al., 2017; Macé et al., 2018). This technology measures changes in the Cerebral Blood Volume (CBV) within the micro-vascularization by Ultrafast Doppler (Bercoff et al., 2011) and spatiotemporal clutter filtering based on Singular Value Decomposition (Demené et al., 2015). Therefore, as in fMRI, it relies on the neurovascular coupling of brain activities.

If the first reports were performed on anesthetized animals (Macé et al., 2011; Osmanski et al., 2014a, 2014b; Urban et al., 2014), more recent studies have involved awake rodents (Macé et al., 2018; Sieu et al., 2015; Tiran et al., 2017; Urban et al., 2015) and awake non human primates during tasks (Dizeux et al., 2019). Recent studies demonstrated the capability of fUS imaging for high resolution mapping of the 3D tonotopic organization in the auditive cortex and deeper structures such as the inferior colliculus in awake ferrets (Bimbard et al., 2018), in neonates through the transfontanellar window (Demene et al., 2017) and in adults in peroperative setting during tumor resection (Imbault et al., 2017). In the present study, our aim was to evaluate the reliability of the technique for mapping the functional organization of the visual cortex in awake primates such as retinotopic maps and ocular dominance columns. Our results in NHP illustrate that the fUS spatiotemporal resolution and sensitivity in deep tissues offers access to activities in non-optically accessible deep cortical layers of the primary visual cortex as well as in even deeper cerebral areas along the calcarine and lunate sulci.

## Results

### Ten 0.5s long stimulations are sufficient to accurately map a cortical activation

We recorded the variations of Cerebral Blood Volume (CBV) using functional ultrasound imaging in two monkeys while they were performing a passive fixation task (Figure 1A). The recording chambers on the two animals were positioned to have two fUS images in contiguous sagittal plans on the visual cortex (ML +7) (Figure 1B and 1C). For monkey S, the recording chamber was positioned just above the calcarine sulcus (dashed line, Figure 1C) to maximize the surface and depth of imaged V1 (Daniel and Whitteridge, 1961; Huff and Dulebohn, 2018; Wandell et al., 2007) (Figure 1C). For monkey T, the recording chamber was positioned above the lunate sulcus to image the contiguous visual areas, V1, V2 and V3 in the same plane (Gattass et al., 1981; Wandell et al., 2007; Zeki, 1978) (Figure 1C). The fUS image was 10 mm in depth and 14 mm in width. On these anatomical fUS acquisitions, we can distinguish cerebral blood fluctuations within the blood vessels that appeared in white in these images. This image enabled us to distinguish the anatomical sulci (calcarine sulcus and lunate sulcus) due to the presence of blood vessels as well as the deep micro-vascularization through the different cortical layers (white stripes).

**Figure 1:**
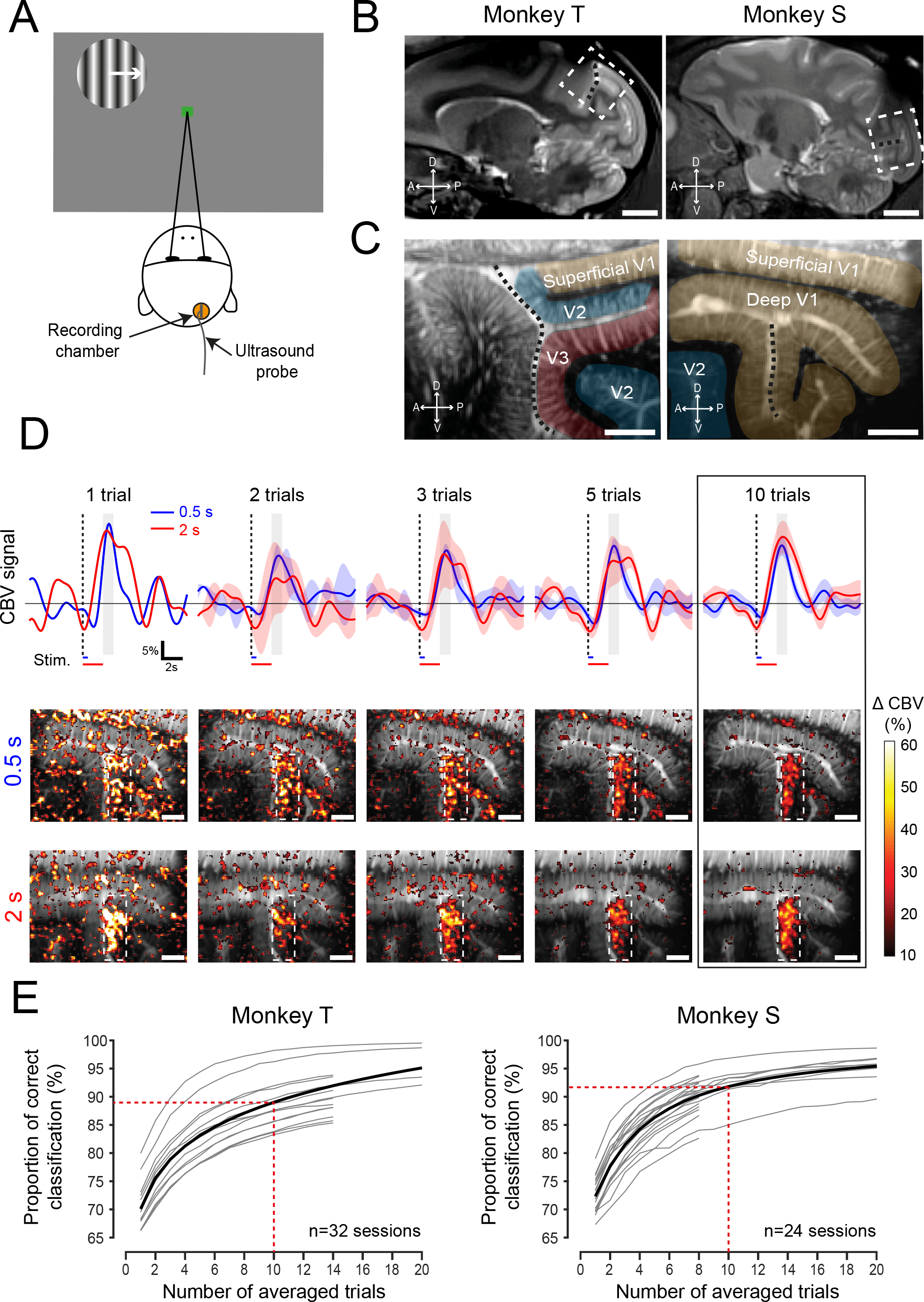
Optimization of the stimulation duration and the number of repetition for fUS imaging. A) Passive fixation task. While fixating a central green square, a peripheral visual stimulus was flashed for 0.5 or 2s. The recording chamber was positioned above the right visual cortex in both animals. B) Anatomical MRI (ML+7mm) showing the localization of the fUS imaging plans (white dotted rectangles) for monkey T (Left) and monkey S (Right). Lunate and Calcarine sulci are respectively highlighted for monkey T (Left) and S (Right) with black dotted line. C) Deep anatomical fUS acquisition. Black and white image represents the amplitude of the mean CBV. The different visual areas are represented by colors (yellow: V1; blue: V2; red: V3). D) CBV measurements during repeated visual stimulation. Top: mean CBV signal variations within the ROI represented in bottom maps by the white dotted rectangles with an increasing number of trials for the two stimulation durations (blue: 0.5; red: 2s, Shaded bars: SEM). Bottom: Activation maps obtained for 0.5s (top) or 2s (bottom) of stimulation for the number of averaged trials (1, 2, 3, 5 and 10 trials). E) Statistical analysis of the measurements showing the proportion of pixel correctly classified with respect to an image generated with 20 other trials (32 sessions for monkey T; 24 sessions for monkey S) with 0.5s stimulation duration. Grey lines represent the evolution for each individual session (Sessions composed by only 1 trial are not plotted). Black lines represent the Naka Rushton fitting result for all the sessions. Scale bars represent 1cm in (B), 2mm in (C) and (D).

We first examined whether CBV variations were modulated by the presentation of visual stimuli. CBV responses were measured in response to 3 different stimuli conditions: 1) when visual stimuli were presented for a short period (0.5s), 2) for a longer period (2s) and 3) when no visual stimuli were presented during the fixation period (Figure S1). For this first test, the visual stimulus was presented at the upper left of the visual field (circle of 2 Degrees of Visual Angles (DVA) diameter centered in [azimuth: −1 DVA; elevation: −10 DVA]). After just a single trial, the CBV increased significantly only within the posterior bank of the calcarine sulcus (Figure 1D). The magnitude and the localizations of these responses were similar for a 0.5s and a 2s stimulation. Such a CBV variation was not observed when the visual stimulus was not presented during the fixation period lasting for 2s (Figure S1).

We then analyzed two important parameters for imaging studies: the influences of the stimulation duration on the recorded hemodynamic responses and the number of needed trials to obtain a clear activation map. We plotted the mean CBV signal within the responsive ROI (white dotted rectangle in activation maps, size: 50×18pixels) according to the period of stimulation (blue: 0.5s; red: 2s). When only ten trials were averaged (Figure 1D, black rectangle), the CBV peak occurs after 2.5s of the stimulation onset for both 0.5s and 2s of stimulation duration. Hence the mean CBV variation was averaged during 1s centered on 2.5s (see grey rectangle) after the visual stimulus onset to compute all the activation maps in this study. The amplitudes of average responses (for n=10 trials) are similar and reached 16.27% ±2.81% for a 0.5s stimulation and 18.5% ±4.65% for a 2s stimulation. Moreover, the Signals to Noise Ratio (SNR) on activation maps (Figure 1D) were in the same range for the short (0.5s) and the long stimulations (2s) (2.78dB and 3.15dB for the respective 0.5s and 2s activation maps). Considering all these results on the two stimulation durations, we decided to use the shortest stimulation duration (0.5s) in the following protocols described in this study.

To reveal responsive areas with functional imaging techniques, another important feature is to define the number of trials, which should be averaged to create a relevant activation map. With only a 0.5s single trial, the responsive region is clearly detected on the activation map with a response peak reaching 25.3% although this map looks quite noisy (Figure 1D). To investigate the influence of averaging on the response reliability, we generated combinations of activation maps computed with different number of trials (1 to 20) comparing each of them with an activation map computed with 20 other trials. The comparison was done by counting the pixel correctly classified, hence, the true positive ones and the true negatives ones. With only 10 averaged repetitions we reached 89.02% of correct classification for monkey S across 24 sessions and 91.81% for monkey T across 32 sessions (Figure 1E). In summary, we obtained accurate activation maps after 10 trials even with a 0.5s presentation of the visual stimulus. All the following measurements were therefore acquired with this short stimulation duration and with at least 10 repeated trials.

### Retinotopic mapping in the whole depth with a high spatial resolution

Considering the promising liability of the results on very few trials, we tested whether fUS imaging could be accurate enough to functionally map the cortex in awake and behaving NHP within a single acquisition session. The different regions of the primate visual cortex are classically defined by their visual sensitivities to a specific eccentricity and angular position in the visual field (Tootell et al., 1998). We therefore tested whether shifting the stimulation locus across the visual field would shift the CBV variations across the visual cortex. The CBV variations were first measured in response to 9 different visual stimulation eccentricities restricted to a half ring in the left visual field. The range of the maximal cortical CBV responses was between 10% and 40% as expected (Figure S2A). Figure 2A presents four different activation maps when visual stimuli were presented respectively close to the fovea (top left: 4-5.5 DVA), at some mid-eccentricities (top right: 8.5-10DVA; bottom left: 11.5-13 DVA) and a larger eccentricity (bottom right: 13-15 DVA). All these CBV activation maps were different for the different visual field locations. To quantify these differences, we selected 10 different pixels in this example along the activation maps (Figure 2B). When the visual stimulation was located close to the foveal region (4-5.5 DVA), CBV increased significantly for pixels #1 to #6 compared to CBV responses in pixels #7 to #10 that increased for more eccentric stimulations. This eccentricity selectivity is generalized for all pixels in figure S2B. This map represents the eccentricities inducing the highest measured CBV responses. This observation is consistent with previous studies showing that superficial V1 processes visual information for low eccentricities unlike deep V1. Moreover, some regions seemed less selective with a large bandwidth (pixel #10 for example) compared to others with a sharp bandwidth (pixel #7 for example). This selectivity disparity is highlighted in Figure S2C representing the bandwidth of the Gaussian fitting for each pixel.

**Figure 2:**
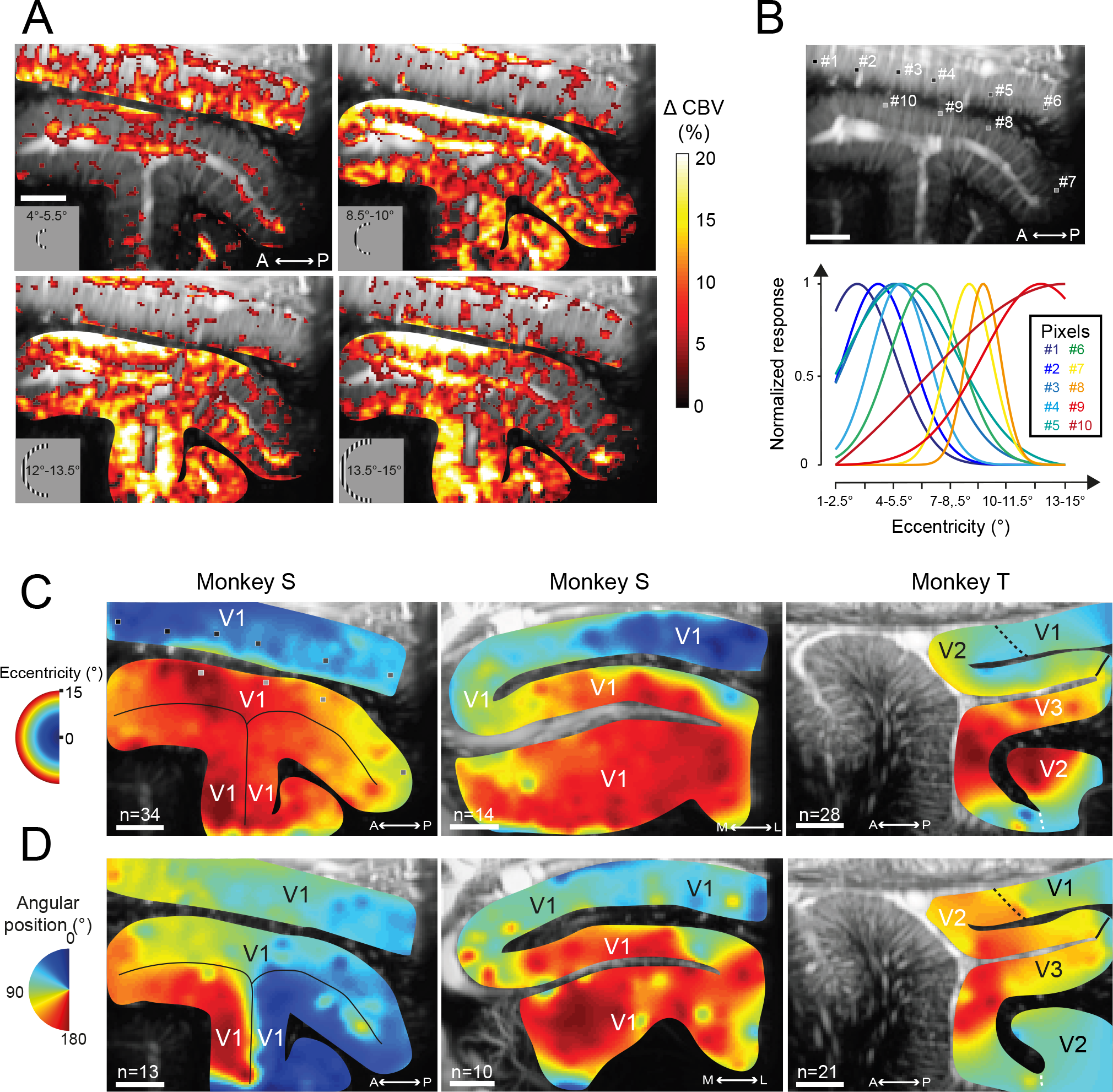
Retinotopic maps of the visual cortex with fUS imaging. A) Activation maps obtained for 4 different stimulation eccentricities in V1 of monkey S (34 averaged trials). B) Evolution of the normalized CBV responses for the 10 chosen pixels as a function of the stimulus eccentricity. C-D) Eccentricity (C) and angular (D) retinotopic maps reconstructed from activations maps as in (A) obtained in a single session for each retinotopic map in the sagittal and coronal plans from monkey S (Left and middle) and monkey T. 34, 14 and 28 trials were averaged to compute respectively the left to right eccentricity maps. 13, 10 and 21 trials were averaged to compute respectively the left to right angular maps. Scale bars in A-D represent 2 mm.

Figures 2C-D illustrate all cortical retinotopic maps for the two monkeys. These retinotopic maps were established with only one recording session for each map. For instance, the first eccentric map was generated with 34 correct trials per condition resulting in an acquisition session lasting shorter than 1 hour (Figure 2C, left). On this map, we observe a consistent representation of the visual field with larger eccentricities represented through the calcarine sulcus in depth and the smaller eccentricities within superficial V1 (Figure 2C, left). The fact that the eccentricity selectivity doesn't change within superficial V1 and within deep V1 with the sagittal imaging plane was expected considering the organization of the visual cortex (Figure 2C left and right). Moreover, the transversal plane confirms the progressive evolution of the eccentricity projection of V1 within the medial fold (Figure 2C, middle) with a foveal representation at the surface whereas deep V1 is more activated by more eccentric locations.

On the angular retinotopic maps, the sagittal plane in monkey S (Figure 2D Left) illustrates that visual stimuli located in the upper visual field induce significant CBV variations in the posterior bank of V1 calcarine sulcus (represented in blue; angular position around 0°). By contrast, visual stimuli presented in the lower visual field induce CBV changes in the anterior bank of V1 calcarine sulcus (represented in red; angular position around 180°). The fact that the angular selectivity doesn't change within superficial V1 with the transversal imaging plane was expected considering the organization of the visual cortex (Figure 2C, middle panel). On monkey T, the V1/V2 and V2/V3 borders have been revealed with the respective vertical and horizontal meridians projection (black dashed and black continuous lines). As expected within dorsal visual areas (V2 and V3d), selective responses correspond to visual stimulations in the lower visual field (represented in orange and red: polar angle between 90° and 180°).

As indicated above, all these retinotopic maps were established after only one acquisition (about 1h of experiment). To illustrate the reliability of the measurements, Figure S2E-F presents two retinotopic maps generated in two different sessions on two different days. Note that the functional and anatomical maps were not fully identical because the position of the probe could not be secured very precisely at the same position each day. However, despite this problem, 77.84% and 89.10% of the pixels had the same eccentric value (±2DVA) for the respective Figure S2E and Figure S2F suggesting that across different behavioral sessions, we could obtain similar retinotopic maps. Together, all these results indicated that functional ultrasound imaging system can be used to accurately map visual responses within different visual areas even within the depth of the visual cortex.

### fUS maps Ocular Dominance bands and reveals cortical layer selectivity of the visual cortex

To further assess the spatial functional resolution of fUS imaging, we examined whether ocular dominance columns (ODCs) can be resolved with this new imaging technology. ODCs have indeed thickness of about 400-700μm in macaque visual cortex (Horton and Hocking, 1996; Hubel and Wiesel, 1972; Levay and Connoly, 1985). To perform this functional map, we measured CBV variations after the presentation of a visual stimulus in the central visual field (limited to 15 DVA) consecutively when one eye (ipsilateral or contralateral) was obstructed (Figure 3A, Left). A map revealing ocular dominance columns (Figure 3A, right) was obtained by subtracting contralateral to ipsilateral evoked maps. In both animals, we found the presence of alternative complementary bands on the superficial cortex. Indeed, OD columns were present in superficial V1 (ROI #1), but also through the calcarine sulcus (ROIs #2 and #3). In contrast, we did not notice such OD columns at the roof of the calcarine sulcus. As visualization of ocular dominance columns is expected if the imaging plane is oriented perpendicular to OD bands observed previously by autoradiography (Hubel and Wiesel, 1972), we performed observations in the same area but in the transversal plane (Figure 3B, Top). In this transversal plane, the OD columns became apparent at the roof of the calcarine sulcus (ROI #4), but they disappeared in the superficial V1 or in deep V1 (ROIs #1, #2 and #3). The presence of ocular dominance columns with fUS technique was confirmed on Monkey T (Figure 3B, bottom) along the superficial V1 (ROI #5) even if they are less distinguishable than on monkey S. Finally, the V1/V2 border on monkey T was revealed with the presence of OD stripes only in V1 (black dotted line). Indeed, for both monkeys, no OD columns were visible in V2 (ROI #6 for monkey S and ROI #7 for monkey S). The reliability of these OD columns was tested by computing the map after shuffling ipsilateral with contralateral blocks (Figure S3A). In this analysis, disappearance of the OD columns confirmed that these are not artefacts due to one acquisition (ipsilateral or contralateral). Indeed, both activation maps (ipsilateral and contralateral) are clearly complementary in response, proving the robustness of this functional map. Moreover, without the manuel cropping (Figure S3A, left), we can notice that OD columns are only visible within V1 and are perpendicularly oriented to the cortical surface.

**Figure 3:**
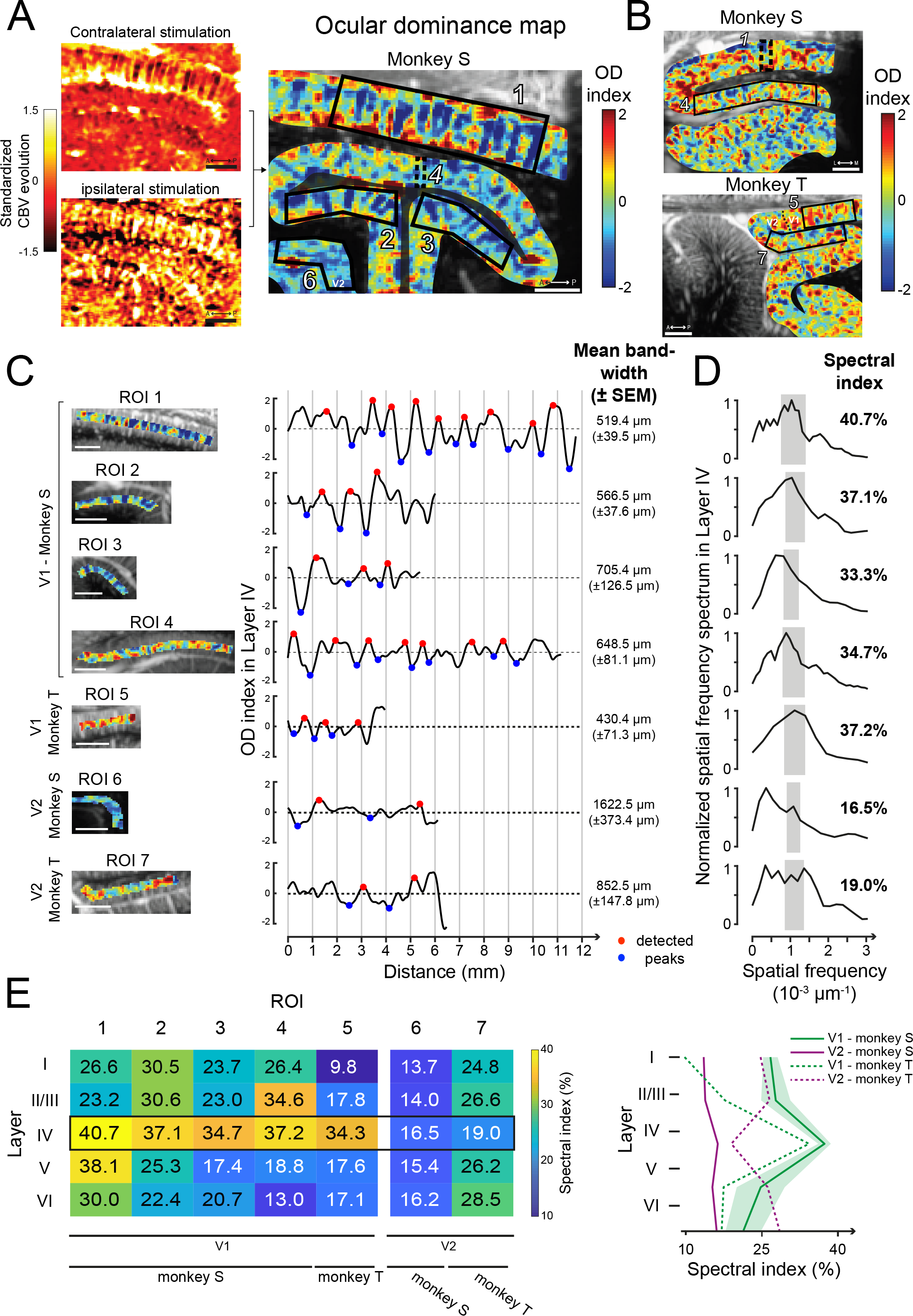
Ocular dominance maps in the visual cortex. A) OD map (right) obtained from the Standardized CBV evolution in V1 for Contra (Top) and ipsilateral (Bottom) stimulations in monkey S (Left). B) OD maps obtained for monkey S in a transversal imaging plane for monkey T in the sagittal plane. Black dotted line: V1/V2 border. C) Mean OD index across layer IV for each ROI (left, black polygons in A-B) showing peaks (red and blue circles) used to compute the mean bandwidth. D) Spatial frequency spectrum of the layer IV for each ROI. Spectral index is computed integrating the spectrum in a bandwidth corresponding at a 350-700μm OD bandwidth range (grey rectangles). E) Table and curves with spectral index computed and averaged for all layers and ROIs (shaded bar= SEM). Scale bars represent 2mm in A-C.

We then wanted to confirm that the bands we observed presented similar parameters than the ones previously described in the literature. To estimate the OD band widths, we selected 7 different ROIs in superficial V1 (#1 and #5), deep V1 (#2, #3 and #4) and V2 (control, #6 and #7). For these ROI, the OD index was plotted along layer IV after having segmented V1 visual cortex according to the Hässler’s scheme (Figure 3C, right panel; see Figure S3B for layer segmentation (Balaram and Kaas, 2014)) (Fig.3C, left panel). Periodic OD index oscillations were clearly visible for ROI#1 and the mean OD bandwidth for superficial V1 is 519.4μm (±39.5μm). This bandwidth was similar within the dorsal and ventral banks (ROI #2: 566.5μm ±37.6μm; ROI#3: 705.4μm ±126.5μm), with those of the roof of the calcarine sulcus (ROI #4: 648.5μm ±81.1μm) and of the superficial V1 of monkey T (ROI#5: 430.4μm ±71.3μm). These measures were in great accordance with the classical range of 400-700μm classically reported for ocular dominance band thickness (Horton and Hocking, 1996; Hubel and Wiesel, 1972; Levay and Connoly, 1985). OD index along V2 did not show the same range of bandwidth (ROI #6: 1622.5μm ±373.4μm; ROI #7: 852.5μm ±147.8μm). Spatial frequency spectra of the OD index in layer IV further confirmed the presence of a periodicity corresponding at the classical OD bandwidth for V1 ROIs (#1 to #5) (Figure 3D). Indeed, a peak was clearly visible around the corresponding 350-700μm bandwidth for these V1 ROIs (a period of two bands is represented by the gray rectangles: 7.1429.10^−4^μm^−1^ < SF < 1.4.10^−3^μm^−1^). By contrast for V2, the selectivity peak was present at a very low spatial frequency in ROI #6 (SF=1.631.10^−4^μm^−1^) and the spectrum covered a large selectivity bandwidth for ROI #7 (0 μm^−1^ < SF < 1.386.10^−3^μm^−1^). To quantify these spectra within the theoretical OD bandwidth, we computed the “Spectral index” as the proportion of the spectrum covering the 350-700μm bandwidth (Figure 3, grey rectangles). This spectral index was superior to 30% for all V1 ROIs when it was lower than 19% for all V2 ROIs. This result was consistent with the notion that such oscillations at the classic OD bandwidths were only present in V1.

The precise spatial resolution of this innovative imaging technique allows us to study the cortical layer segmentation. OD bands were mainly reported in cortical layer IVc in visual area V1 and interpreted as a layer specificity of thalamic projections (Hubel and Wiesel, 1972; Levay and Connoly, 1985). To further assess this layer selectivity, we computed the normalized spatial spectrum for all layers of each ROI to extract all spectral indexes (Figure 3E). For V1 ROIs (#1 to #5), this analysis revealed clear increases of the spectral indexes in layer IV compared to layer I, II/III, V and VI. These peaks in layer IV were similar for both monkeys in V1 whereas there was no such an increase through layers in V2 (Figure 3E, right panel). These observations are in accordance with the first observations of Hubel & Wiesel (Hubel and Wiesel, 1968, 1972). However, even if the spectral indexes were always in the highest range for layer IV (34.3% to 40.7%), some ROIs also showed high values within layer II/III (ROIs #2 and #4, respectively 30.6% and 34.6%) and within layer V (ROIs #1: 38.1%). This study indicates that fUS imaging can be used to measure OD columns and that OD columns may extend to layer II/III and layer V of the V1 visual cortex.

## Discussion

A key aim of this study was to test whether we could record and localize neuronal activity within the visual cortex of awake non-human primates at a mesoscale (ranging from 100 microns to 2 centimeters) using an innovative imaging technique: functional ultrafast ultrasound imaging (fUS imaging). In response to different mapping protocols, we have revealed precise retinotopic maps consistent with validated neurofunctional knowledge of primary visual cortex of NHP. This range of spatial resolution and the field of view in depth enabled us to distinguish ocular dominance selectivity of cortical layers in the whole V1. fUS imaging overcomes some disadvantages of conventional techniques (fMRI, optical imaging methods) for different reasons we detailed bellow.

### The first deep functional imaging mapping through all cortical layers of the visual system

With the OD mapping with fUS imaging we demonstrated a clear advantage of imaging activity *in vivo* with a high spatiotemporal resolution in depth. Indeed, since Hubel & Wiesel have demonstrated the existence of OD cells in V1 with electrophysiology recordings (Hubel and Wiesel, 1968), most of the following studies have used autoradiographic methods needing the sacrifice of the animals (Levay and Connoly, 1985; Tootell et al., 1988; Wiesel et al., 1974). In functional imaging, OD bands were also detected by optical imaging methods like Intrinsic-signal Optical imaging (Kaskan et al., 2007), VSDi (Blasdel and Salama, 1986) or calcium imaging (Nauhaus et al., 2016) but this demonstration was always limited to the superficial cortical layers because optical method cannot reach deep layers and deep cortex. Here we showed that fUS imaging overpasses this limitation mapping the OD columns *in vivo*, in the whole cortex depth. Ocular dominance columns were classically described in layer IV of V1 (Levay and Connoly, 1985) as confirmed by the recording of cells responding to the stimulation of one eye (Hubel and Wiesel, 1968).

Our observation of an extension of OD columns in layer II/III and layer V is consistent with previous anatomical studies showing some continuities between OD columns in layer IV and such OD columns in layer V or cytochrome oxidase blobs in layer II/III as indicated by autoradiography or immediate-early genes immunolabelling (Takahata et al., 2009; Tootell et al., 1988). However, in the visual cortex, large blood vessels are extending vertically. As a consequence, one may wonder whether a vertical alignment of large blood vessels in V1 could artificially extend the columns in layers II/III and V. Indeed, the blood circulating in a closed system generates a peripheral influence on the measured signal (Carandini et al., 2015; Uhlirova et al., 2016; Vanzetta and Grinvald, 2008). Although it is not possible to completely refute this hypothesis, it is not probable here. Indeed, our analysis of the ultrafast ultrasound technology filters the signal produced by arterioles, capillaries and venules based on their blood velocity (between 1 and 25mm/s) and discriminate it from the influence of larger vertical vessels (Demené et al., 2016). Moreover, Rungta et al. recently measured the red blood cells velocity variations across the different vascular compartments (from the juxta-synaptic capillary backward to the feeding pial arteriole) in the mouse olfactory bulb, in response to odour. Despite an heterogeneity of blood velocity increases according to the vessel type, the largest proportional increase of this hemodynamic parameter was observed in the juxta-synaptic capillaries (Rungta et al., 2018), providing evidence for the very high sensitivity of measuring CBV evolution to accurately localize neural activations. Furthermore, with the dynamic filtering out of large vessels in fUS imaging, the smaller change in blood velocities may explain why no spatial blurring of the functional response was reported in recent studies distinguishing small cortical areas with fUS imaging (Bimbard et al., 2018; Macé et al., 2018; Osmanski et al., 2014a). For instance, revealing the tonopic map in awake ferret with fUS imaging, Bimbard et al. were able to discriminate pixels spaced by 300μm with their tuning curves. Such a resolution is compatible with layers and OD columns discrimination in the visual cortex of NHP. In fact, different anatomical studies have also suggested some continuities between OD columns in layer IV and such columns in layer V or cytochrome oxidase blobs in layer II/III as indicated by staining by autoradiographic markers or immediate-early genes methods (Takahata et al., 2009; Tootell et al., 1988). Therefore, our observation of the ocular dominance columns from layer II/III to V provides the first *in vivo* evidence of this V1 organization with functional imaging.

The weak distinction of the OD columns in deep cortex of V1 could be due to smaller column sizes below the resolution limit of the technology. Indeed, OD bands are thinner for high eccentricities (Adams et al., 2007; Horton and Hocking, 1996; Van Essen et al., 1984). OD bands parallel to the ultrasound probe could also provide an alternative explanation. This could likely be the case for the OD columns at the roof of the calcarine sulcus as indicated by a previous study (Levay and Connoly, 1985). Designing a new probe with an higher US frequency (30MHz) would increase the spatial resolution for such measurements by a factor 2 (Gesnik, 2017). Such an increase in spatial resolution could enable investigators to identify other functional structures like orientation selectivity columns in V1, or CO-Blobs although the maximal imaging depth would be reduced (Blasdel, 1992; Blasdel and Salama, 1986; Grinvald et al., 1986; Ikezoe et al., 2013; Nauhaus et al., 2016).

### A fast, easy and accurate mapping

Showing that only ten short stimulations (0.5s) are enough to accurately map a cortical activity proved a big interest of this method compared to fMRI which is less sensitive and more time-consuming even the last laminar fMRI technique (review: Lawrence et al., 2017) using Ultra High Field MRI to study visual cortex in human (Dumoulin et al., 2017). Indeed, mapping the anatomic micro-vascularization with this ultrasound system is very fast: 1 second is enough to capture one imaging plane with a spatial resolution of 100×100×400 μm^3^ and a FOV of 1.4×2 cm^2^ compared with a 7T MRI which typically requires couple of minutes to provide 0.5mm isotropic acquisition of the brain (Zwanenburg et al., 2011). Moreover, MRI acquisitions can be stressful for monkey and there are different electromagnetic constraints (Tsao et al., 2008; Vanduffel et al., 2001; Yue et al., 2013), whereas fUS imaging can be easily performed on awake behaving monkey. Therefore, fUS imaging can thus easily be exploited for anatomical acquisitions in recording chambers to guide injections or electrode implantations, especially in depth.

However, some functional structures are anatomically oriented but fUS only records one 400μm thick imaging plane oriented in depth. Consequently, it would be tricky to image functional structures when their orientation - or position - are parallel to the imaging plane. This spatial restriction could be compared to the deep axis limitation with optical imaging. The solution is, when possible, to perpendicularly rotate or shift the probe within the recording chamber, to image the cortex with a different angle, allowing thereby to solve the spatial problem. A 3D reconstruction could thus be made like previously achieved with parallel imaging planes (Demené et al., 2016; Gesnik et al., 2017; Macé et al., 2018).

Given the good SNR of this method, fUS imaging could quickly become a gold standard to functionally map a region for further electrophysiological studies, DBS implantation, or localized drug delivery. Indeed, electrophysiological methods are too time-consuming for such preliminary mapping (Daniel and Whitteridge, 1961; Dow et al., 1981; Hubel and Wiesel, 1974; Talbot and Marshall, 1941; Van Essen et al., 1984) whereas fUS imaging can generate a retinotopic map with only one-hour session. Furthermore, considering a perfect session with only correct fixations for all trials, only 420 seconds of acquisition is needed to reconstruct the polar angle map with fUS imaging when it takes 3200 seconds (more than 7 times) with fMRI (Arcaro and Livingstone, 2017). Moreover, fUS imaging has a better spatial resolution allowing mapping the OD bands in non-human primates when it’s currently unreachable with fMRI.

### A slightly invasive and long-term imaging technique

Although this fUS imaging technique requires a craniotomy for primate studies - but not necessary for rodents (Errico et al., 2016; Tiran et al., 2017) - the dura is kept intact. Indeed, ultrasonic waves propagate without major reflections or diffraction effects through soft tissues. This property means that dura, coagulated blood or others optical barriers do not strongly affect the SNR. Within several months, we observed a slight decrease of the SNR, which may be due to the dura thickening but it remained negligible for our needs. Therefore, fUS imaging provides a powerful imaging technique for long term chronical studies. Its field of application is adapted to map superficial and deep cortical areas (up to 2cm deep). Increasing further the depth of acquisition up to subcortical areas like the thalamus for example would require reducing the US frequency and thus decrease the spatial resolution. Using a 6Mhz frequency probe, we were able to image throughout the macaque brain (Dizeux et al., 2019).

### A continuum of brain imaging technology

Brain imaging is becoming a major field of technological innovation to investigate brain function in normal and pathological conditions. The resolution of fMRI is increasing further by increasing the power of the magnets. However, such magnets are difficult to bring into surgical rooms and in laboratories. Recently, magnetoencephalography was adapted to generate measurements in freely moving patients with a very high temporal resolution but a low spatial resolution (Boto et al., 2018). By contrast, we show here that fUS imaging provides a very good spatial resolution at the mesoscale level through the cortical depth as long as a window can be achieved on the skull. This new technology should therefore fill a major gap in brain mapping within surgical rooms and laboratories when optical techniques are not relevant due to depth concerns. Although yet limited, fUS imaging technology has already been demonstrated on human babies during transfontanellar imaging (Demene et al., 2017) or on human patients for peroperative imaging during brain surgery (Imbault et al., 2017).

## Materials and Methods

### Experimental apparatus

We collected data from two Rhesus monkeys (Macaca mulatta, “S” a male aged 13 years and “T” a female aged 11 years), weighting 13 and 9 kg respectively. These monkeys were individually housed and handled in strict accordance with the recommendations of the Weatherall Report about good animal practice. All experiments were conducted after validation of the European Council Directive (2010/63/EU) and the study was approved by the French ministry and the institutional and regional committees for animal care (Committee C. Darwin, registration number 9013).

Under isoflurane and aseptic conditions, we surgically implanted titanium head-posts and recording chambers according to the surgical procedure described in a previous study (Valero-Cabre et al., 2012). Briefly, monkeys were deeply anesthetized with an IM injection of blend of Ketamine (0.3mg/Kg) and Dexmedetomidine (0,015mg/kg) for initial sedation and maintained with isoflurane gas (1-2%) during the surgery. Heart rate, temperature, respiration and Sp02 are monitored. Pain medication was given prior the surgery with butorphanol (0.2mg/kg i.m.) when the animal was being mechanically ventilated (Hallowell EMC model 2000) and buprenorphine (0.015mg/kg i.m) routinely given at the end of the surgery (and for reverse butorphanol effects). After surgery, an antibiotic was also administrated to avoid infections (Clamoxyl LA 15mg/Kg or Duphacycline LA 20mg/Kg). The head post was positioned on the medial line and was rostral enough to allow the subsequent implantation of a recording chamber above primary visual area. We waited a recovery period of 6 weeks before fixating the animal's head.

After the recovery period, animals were trained to perform a passive fixation task (Figure 1A). When their performance reached a significant threshold, a recording chamber was implanted above the visual cortex (Figure 1B) based on some MRI scans (3T MRI; T2* sequence; 0.5mm isotropic). Monkeys were deeply anesthetized and monitored during the surgery. The recording chamber (Crist Instruments / CILUX chambers 6-IAM-J0) was positioned and centered above the right calcarine sulcus for monkey S (ML +7mm; AP −22mm) and above the right lunate sulcus for monkey T (ML +7mm; AP −10mm; DV +38mm). The dura mater was kept intact during these surgeries.

### Visual stimuli

Each behavioral session lasted a maximum of two hours. The animals were seated in a primate chair (Crist instuments, Inc., Hagestown, MD) with their head fixed and placed in front of a computer screen (Liyama Prolite XB2783HSU; gamma corrected, resolution: 1920 x 1080, running at 60Hz) 53 cm away in a darkened booth. Mean screen luminance was controlled (9 cd/m^2^). Eye position was monitored using an EyeLink 1000 infrared eye-tracking system (SR Research Ltd, Ottawa, Ontario, Canada). Experiments were controlled by EventIDE software (OkazoLab). The monkey started the trial by fixating a central green square (0.2 DVA) for 500ms within a tolerance window of 1 to 1.4 DVA. A visual stimulus on the peripheral location was then presented for 0.5s or 2s. We used drifting sinusoidal gratings as visual stimuli with a fixed temporal frequency (3 cycles/s) and a fixed spatial frequency (1 cycle/deg). The animals were rewarded by a small drop of liquid (water) at the end of each correctly performed trial. We imposed an inter-stimulus interval of 3s. We also used control trials with the same temporal organization but without any peripheral visual stimulus (Figure S1). All conditions (visual and control) were randomly interleaved. We collected in average 20 trials per visual condition.

We obstructed the right eyes of both monkeys with an opaque visor to generate the retinotopic maps of the visual cortex. We focused on recording 3 different functional maps: one based on the eccentricity, one based on the angular positions and one for the ocular dominance columns. For the eccentricity map, we used a set of 9 different visual conditions to eccentricities from 1.5 DVA to 15 DVA. Each stimulus is defined as a hemi-concentric band of 1.5° width centered on the central fixation point. For the angular position maps, we used 12 different visual stimuli: each stimulus was defined by a 15° of angle width, 0° to 180° from 1.5 DVA to 15 DVA of eccentricity. To reveal the ocular dominance columns within the primary visual cortex, either the right or left eye of the monkey was obstructed and we used a “full field” vertical grating (spatial frequency: 1cycle/deg; temporal frequency: 3cycles/s) from 1.5 DVA to 15 DVA of eccentricity.

### fUS acquisition sequences

We recorded fUS images using a linear ultrasound probe (custom design, 128 elements, 15 MHz, 110μm pitch / Vermon, Tours, France) driven by a modified ultrafast ultrasound research scanner (256 electronic channels, 60 MHz sampling rate / Supersonic Imagine; Aix-en-Provence, France). All details from technical procedure and statistics analysis of fUS imaging technique were described in several previous publications from our group (Gesnik et al., 2017; Macé et al., 2011; Osmanski et al., 2014a). Briefly, fUS images were acquired by repeated emissions of a set of 15 planar ultrasonic waves (Pulse Repetition Frequency: 7500 Hz) tilted with different angles ranging from −14 to 14 degrees (Montaldo et al., 2009) at 2° step in order to obtain one high-quality ultrasound image. To sample cerebral blood volumes (CBV) variations, we repeated this sequence 200 times at 500 Hz frame rate-that corresponds to a 400ms acquisition time. In order to remove tissue motion artifacts from the dataset, we applied a recently developed clutter filter technique based on Singular Value Decomposition (Demené et al., 2015). Finally, one ultrasensitive Doppler image was formed for each ultrafast data block with a sampling rate of 1Hz by averaging the 200 compounded and filtered ultrasonic images. Therefore, we extracted cerebral blood volumes (CBV) from ultrasensitive Doppler images with a sampling rate of one image per second. During a behavioral session of 60 minutes, we obtained 3600 images (98×128pixels). We could image 14mm along the cortical surface and 10 mm in depth. Our spatial resolution was 0.11 x 0.1 mm and the width was 0.4mm.

When working on retinotopic maps, we usually only performed all fUS imaging acquisitions required for a complete map during the behavioral session. For ocular dominance maps, we performed all required acquisitions on both eyes during the session.

### Data processing

We then analyzed all the fUS images acquisitions using Matlab (version R2017b; Mathworks). For each behavioral session, a log file is created by EventIDE which was used to synchronize offline CBV signal with the beginning of each trial. CBV signals were extrapolated, 3D smoothed (convolution kernel: [3 3 3]; gaussian filter, SD=0.65) and normalized with a baseline (average of the signal 5s before the onset of the visual stimulus). The peak of CBV response was detected ~2.5s after the onset of the visual stimulus (Figure 1D). The response of a pixel (0.11 x 0.1 x 0.4 mm) was thus quantified as the mean CBV level within 1s period: from 2 to 3 s after visual stimulus onset.

To compute the proportion of correct classification (Figure 1E) we only used sessions composed by at least 20 repetitions of the same visual condition (n=32 for monkey T, n=24 for monkey S). For sessions where the number of repetitions was higher than 40, we split these ones to obtain artificial sessions of 40 trials. We then randomly selected 20 trials within the session to compute a reference activation map and we used a 10% CBV increase threshold to binarize this map. For a considered number of averaged trials, we selected 100 random trials combinations (excluding trials used for reference activation map) to compute the different binary activation maps with the same thresholding (10%). We then measured the mean proportion of correct classification summing only true positive and negative pixels through the whole map compared to the reference one. We repeated this process for each number of averaged trials (1 to 20) and for each session. We used the least squares method to fit a Naka Rushton model on our data.

To reconstruct retinotopic maps, a 3D data matrix is constructed with the different activations maps for each visual condition. A Gaussian fitting is applied for each pixel and the coefficient of determination is computed to threshold (R^2^>0.02) the relevant pixels (colored pixels in Figure S2D). Indeed, only pixels with responses retinotopically modulated, so with a Gaussian behavior, are relevant to reveal retinotopic organization. In order to increase readability of the images, after obtaining the peaking index map (index of the fitted gaussian peak for each pixel, Figure S2B), a 2D median filter, an interpolation and a 2D mean filter (averaging with the 2 neighboring pixels) are applied. However, retinotopic organization is clearly present also before smoothing, (Figure S2B). The map is then manually cropped to only show the responses in cortex (Figure S2E, left).

To reveal the OD map, we computed the two activation maps (ipsilateral and contralateral) and we standardized them. We then subtracted the ipsilateral map from the contralateral map to obtain the OD map with the OD index values. We manually cropped these maps to reveal only the visual cortex. The layer segmentation was performed by indicating manually the top and the bottom of the cortex of the ROI, computing the distance for each pixel between both and indexing them using the Hässler’s scheme (Balaram and Kaas, 2014). The evolution of the OD index across the cortex represents the centered smoothing spline fitting (least squares method) of the datas within the considered layer. The distance corresponds to the distance from the first intracortical column to the far left of the ROI (the far anterior or lateral). The extremum (blue and red dots) were obtained with a function finding the local extrema with a 0.75 OD index minimal peak prominence and an absolute value superior to 0.25 OD index. The mean bandwidths were computed using the averaged half distance between 2 maxima and 2 minima. We then computed the spatial spectrum for each layer of each ROI with a fast Fourier transform. The spatial spectra were filtered with a moving average (with 3 samples) and normalized by their maximum. We defined the spectral index as the proportion of the spectrum covering the 350-700μm bandwidth.

To check the robustness of the OD map, we shuffled the ipsilateral blocks with the contralateral blocks. To do this, we firstly averaged all even blocks (ipsilateral and contralateral) to obtain a first activation map. We then used all residual odd blocks (ipsilateral and contralateral) to compute the second activation map. We then applied the same algorithm as described above to compute this final artificial OD map from these two “even and odd blocks” maps.

---

**Sup Fig1:**
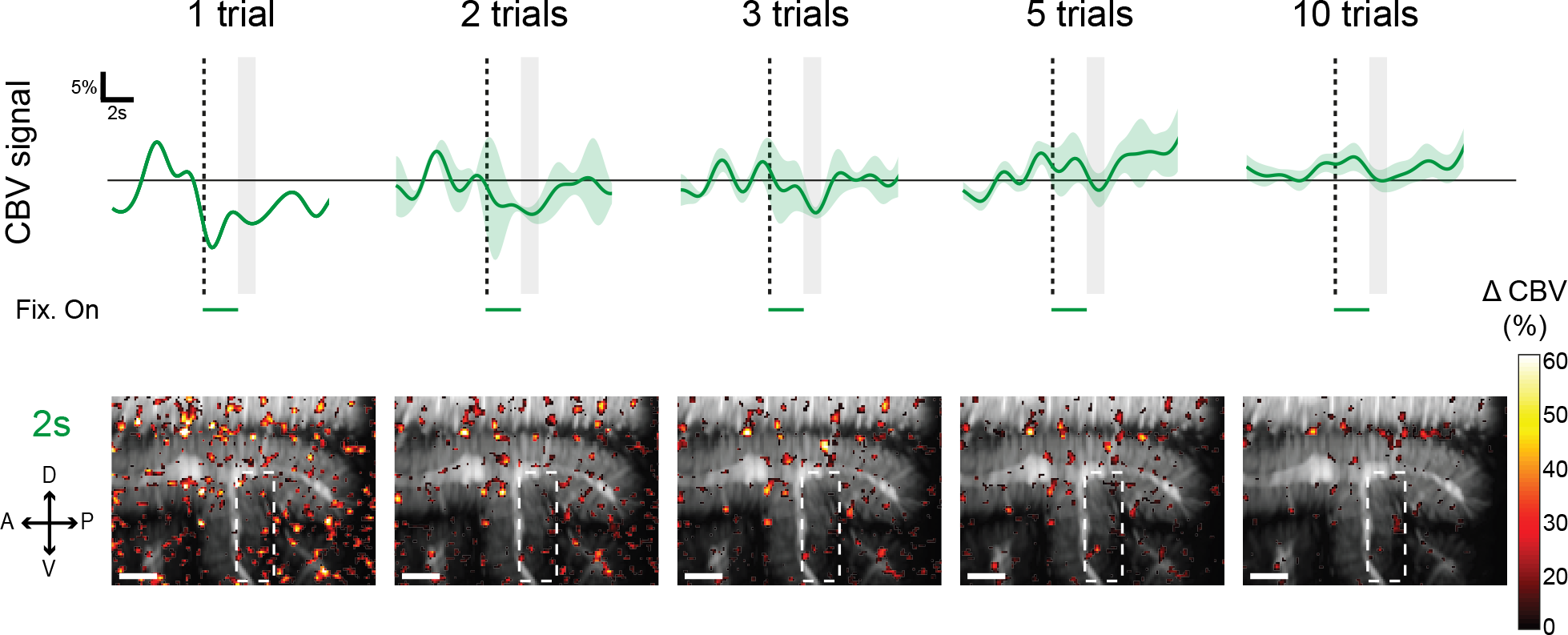
Control acquisition without any visual stimulation during a 2s fixation task. Top: mean CBV signal evolution within the ROI represented in bottom maps by the white dotted rectangles. Shaded area represents the SEM. Bottom: Activation maps obtained according the number of averaged trials. Scale bar represents 2mm.

**Sup Fig2:**
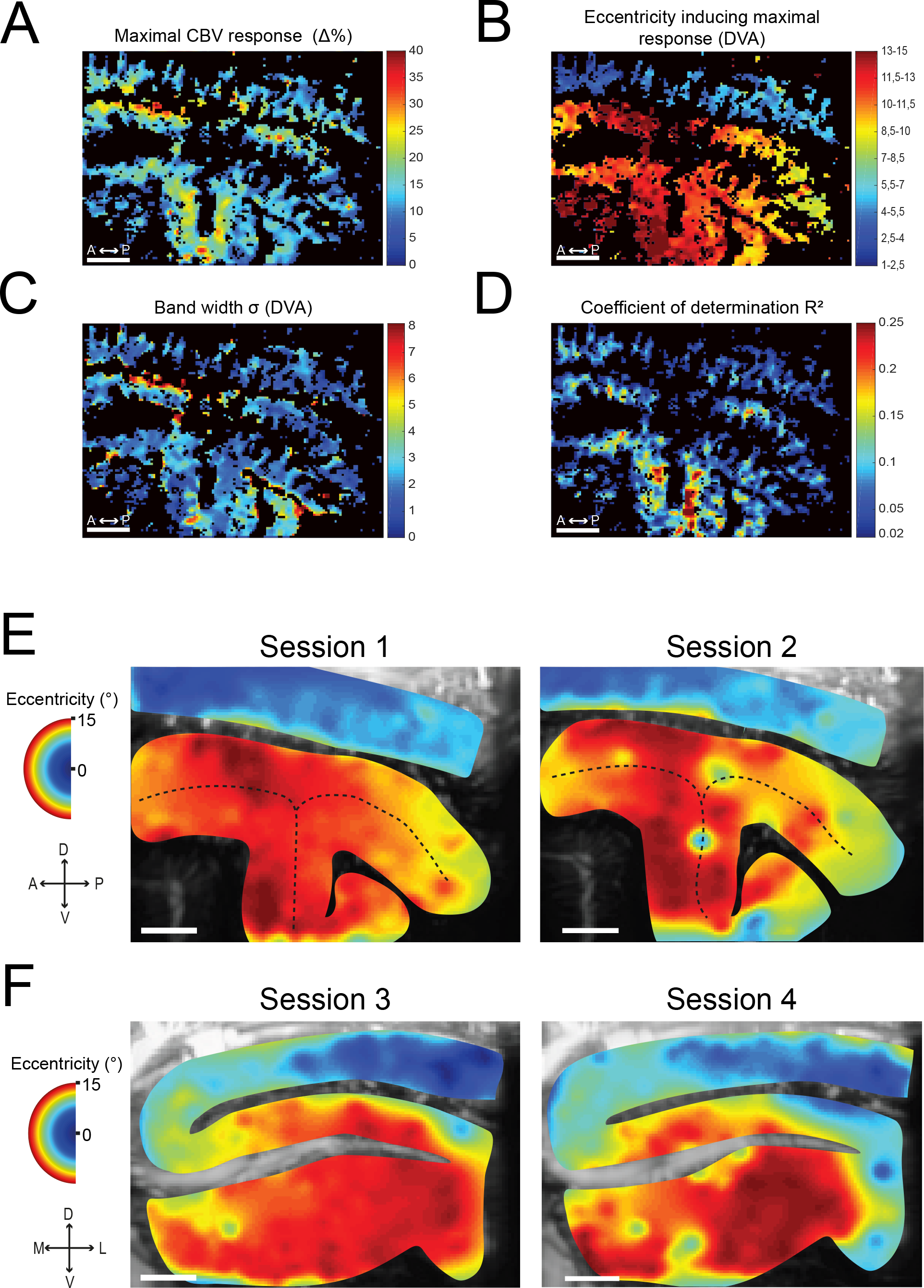
Reproducibility of the fUS imaging retinotopic maps. A-D) Different analyses on activation maps illustrated in Figure 2A. Only significant pixels are shown (threshold: R^2^>0.02). (A) Maximal CBV response map for all eccentricity stimulations. (B) Retinotopic map obtained without extrapolation and smoothing. (C) Selectivity map represented with the Gaussian bandwidth. (D) Coefficients of determination map for a Gaussian curve fitting. E-F) Two eccentricity (E) and two angular (F) retinotopic maps obtained from four different sessions on four different days in monkey S. Note the slight differences in the functional maps and on the anatomical maps that relied on the slight differences in the probe position during the different acquisition day. Scale bars in A-F represent 2 mm.

**Sup Fig3:**
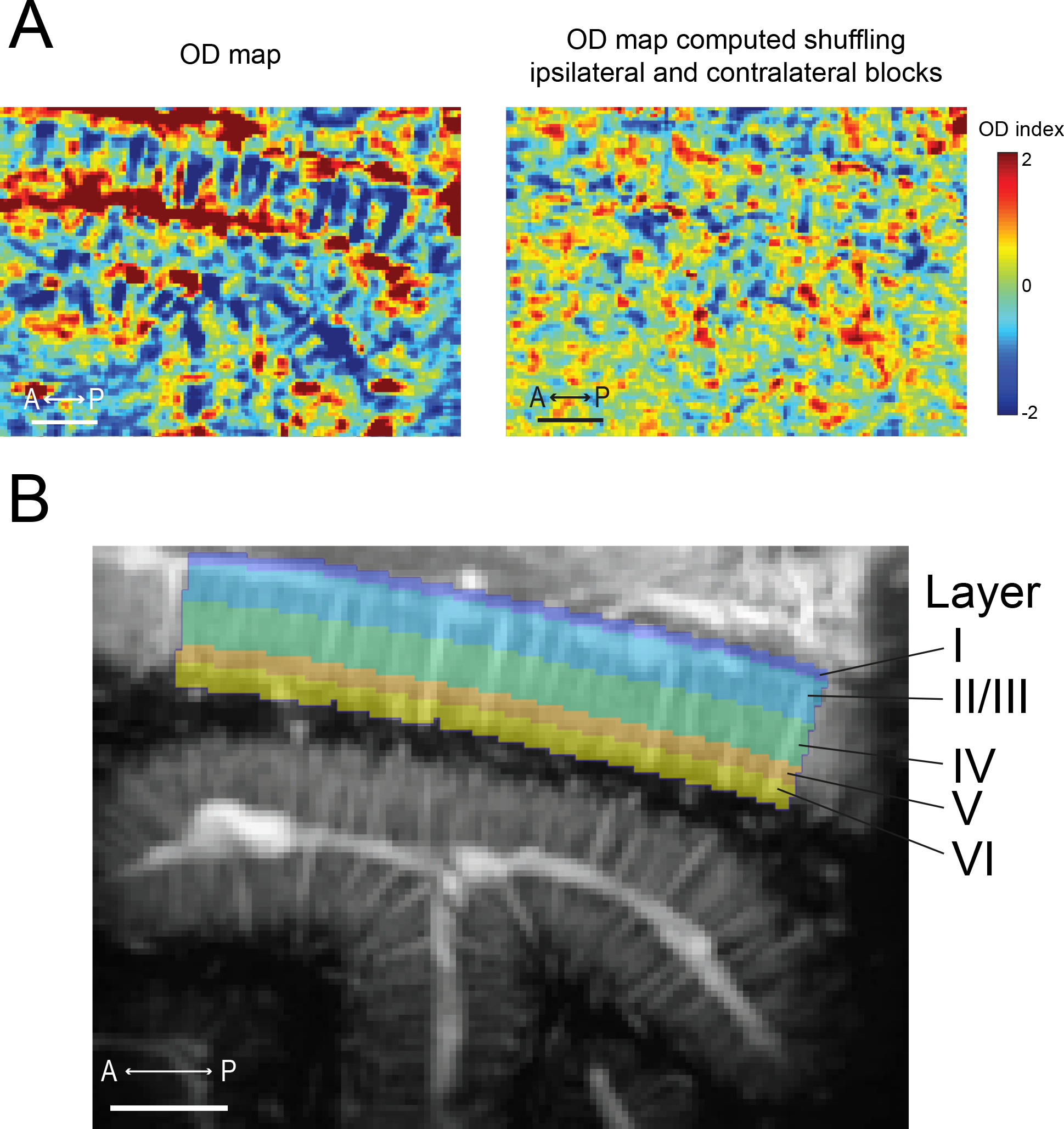
Analyses of OD maps. A) OD map (left) obtained for monkey S without cropping by subtracting ipsilateral blocks from contralateral ones with a subsequent normalization and the corresponding control OD map obtained for the same acquisition but shuffling ipsilateral and contralateral blocks such that even blocks are subtracted to the odd ones prior to normalization. B) Example of layer segmentation computation for ROI #1.

